# Multiple strategies for heat adaptation in rice endosperms revealed by on-site cell-specific analysis

**DOI:** 10.1101/235515

**Authors:** Hiroshi Wada, Yuto Hatakeyama, Yayoi Onda, Hiroshi Nonami, Taiken Nakashima, Rosa Erra-Balsells, Satoshi Morita, Kenzo Hiraoka, Fukuyo Tanaka, Hiroshi Nakano

## Abstract

Plant cells have multiple strategies to adapt to environmental stresses. Rice endosperms form chalkiness in a part of the tissue under heat conditions during the grain-filling stage, although nitrogen supply reduces chalky rice. Air spaces formed in the cells cause an irregular light reflection and create chalkiness, yet what exactly occurs remains unclear at cell level. Through on-site cell-specific analysis, we show that heat-treated cells adjust osmotically and retard protein synthesis to preserve protein storage vacuoles in the cytosol, resulting in air space formation. Application of nitrogen enhances heat tolerance to sustain protein body and amyloplast development during strong osmotic adjustment, which diminishes air spaces to avoid chalkiness. Hence, we conclude that rice endosperm cells could alter organelle compartments spatially during the heat adaptation, depending on the available nitrogen level. Our findings provide new insight into the cellular mechanism of rice chalky formation as a strategy for heat acclimation.

## Introduction

Rice chalkiness is a critical trait that determines the rice quality (Hoshikawa, 1989). Currently, an increase in chalky rice induced by several environmental stresses, such as high temperature during the grain-filling stage, has been frequently observed under global warming (Tashiro and Wardlaw, 1991a, 1991b; Morita et al., 2016). In chalky rice, air spaces formed in a part of endosperms among the loosely-packed starch granules causes significant random light reflection, thereby turning them into chalk (Tashiro and Wardlaw, 1991b). One of the major chalky rice, called ‘white-back rice’, frequently induced by exposure to high temperatures at the early ripening stage, exhibits chalkiness in the outer endosperms longitudinally aligned along the dorsal side of the kernels, where a greater abundance of protein bodies (PBs) are distributed than in the ventral side of the endosperm (Ellis et al., 1987; Hoshikawa, 1989). Moreover, white-back rice is known to be decreased by supplying nitrogen (N) prior to heat conditions (Perez et al., 1996; Wakamatsu et al., 2008). Since chalky formation occur only a part of endosperms, this phenomenon can be a cell-specific event. However, our current understanding of heat-induced rice chalkiness mostly relies on tissue analysis (e.g., Yamakawa et al., 2007); the underlying mechanism in cellular metabolism prior to the air space formation remains unclear. Moreover, the possible role of N during the heat response has not been spatially addressed at the metabolite level.

Storage proteins are major components that accumulate 5–8% in rice endosperms (Juliano, 1985; Hoshikawa, 1989). These proteins are typically stored into two types of protein bodies (PBs), called PBI and PBII (Tanaka et al., 1980; Herman and Larkins, 1999; Shewry and Halford, 2002; Onda, 2013). PBI is a small spherical granule of 1–2 μm in diameter with concentric rings of various electron densities, which is originated from the rough endoplasmic reticulum (rER), and it accumulates prolamins after synthesizing on the ER membrane (Bechtel and Juliano, 1980; Tanaka et al., 1980; Yamagata and Tanaka, 1986; Li et al., 1993; Saito et al., 2012). In contrast to PBI, PBII is a granule with irregular shape, originated from protein storage vacuole (PSV). The diameter of matured PBII typically ranges between 2 to 4 μm, larger than PBI, and glutelin and globulin are both stored in PBII after being synthesized on the rER (Yamagata et al., 1982; Krishnan et al., 1986; Yamagata and Tanaka, 1986; Li et al., 1993). Numerous studies have examined the mutants exhibiting abnormal PB formations (Takemoto et al., 2002; Onda et al., 2009; Wang et al., 2009; Wang et al., 2010; Fukuda et al., 2011; Nagamine et al., 2011; Onda et al., 2011; Ren et al., 2014). Interestingly, these mutations showed chalkiness and are described as “floury endosperm” (Takemoto et al., 2002; Wang et al., 2010; Fukuda et al., 2011; Han et al., 2012; Li et al., 2014; Ren et al., 2014), suggesting that abnormal PB development might be associated with chalky formation in heat-stressed rice. Regarding chalky formation, changes in starch metabolism have received most attention (e.g., Yamakawa et al., 2007; Zhang et al., 2011); however, little has been studied on the possible changes in protein synthesis including PB development. Considering the consistency of the spatial PBs localization of PBs (Ellis et al., 1987; Hoshikawa, 1989) and chalky zone in the outer endosperms of the dorsal side, we hypothesized that protein synthesis might be cell-specifically suppressed under heat conditions, resulting in an inhibition of PB development prior to chalky formation. Some appropriate cell-specific analysis was required to examine our hypothesis.

Recently, single cell metabolomics has been further expanded due to the technical improvement of the mass analyzers (i.e., Orbitrap mass spectrometer) and applied in plant research (Gholipour et al., 2013; Fujii et al., 2015). A cell pressure probe, originally invented by Steudle’s group (Hüsken et al., 1978) and long-used for measuring the cell water status, has been adopted as a picolitre pipette to establish a new type of “analytical method that can be performed in intact plant cells” by combining with an Orbitrap mass spectrometer (Gholipour et al., 2013). More recently, the resolution and sensitivity of the method have been improved by introducing an internal electrode in the capillary holder and adopting the mixture of an ionic solution and the silicone oil in the quartz capillary (Nakashima et al., 2016). This type of cell metabolomics appears to be a robust and powerful method for performing cell-specific analysis. However, most of the cell metabolomics, including this method, have been confined to laboratory use at room temperature. To the best of our knowledge, no attempt has been made to further improve the cell-specific analytical method that allows investigation of metabolic response to some environmental stimuli, such as temperature responses in developing crop plants.

To overcome such limitation at the analysis, we have established a new on-site cell-specific analytical method by combining picolitre pressure-probe-electrospray-ionization mass spectrometry (picoPPESI-MS, referred as ‘IEC-PPESI-MS’ in Nakashima et al., 2016) with an environment control to be used for conducting a cell-specific analysis in the cells growing under heat conditions, which were expected to form chalkiness. Under the temperature equilibrium conditions, this analytical method allows to perform on-site real-time metabolites profiling in the target endosperm cells growing at a set temperature (i.e., 34°C) without any pretreatments (see Nakashima et al., 2016). By combining with a transmission electron microscopy (TEM), it has been shown for the first time that (i) PSVs could be preserved in the cytosol by osmotic adjustment under heat conditions to lead chalky formation as a heat survival strategy and (ii) N-supplied cells increased protein synthesis rate at high temperature to avoid chalkiness.

## Results

### Rice Appearance

In both non-heat (26°C) treatment and N application followed by non-heat (N+26°C) treatment, there were essentially no white-back kernels and the same occurrence of perfect kernels was observed (Table 1). Under heat (34°C) treatment, the proportion of white-back kernels (see 34°C in Figure 1B) increased, reaching to 66.4%, and resulted in a substantial decrease in perfect kernels (Table 1). In contrast, N application followed by heat (N+34°C) treatment ameliorated rice appearance by remarkably decreasing white-back kernels, reaching to 7.6%, but with an approximately 40% increase in perfect kernels (Figure 1C). Moreover, partially ameliorated chalky kernels were classified as other kernels. There was no treatment difference between kernel dry weights in both perfect rice and white-back rice (Table 1), although the average dry weight of pooled kernels declined in 34°C and N+34°C treatments (Table 1). No difference was observed on rice appearance between 26°C and N+26°C treatments and therefore, the following experiments were conducted in 26°C, 34°C, and N+34°C treatments.

**Table 1.** Filled kernels, rice appearance including the occurrence of perfect kernel and white-back kernel, and kernel weight in 26°C, N+26°C, 34°C, and N+34°C treatments.

| Treatment | Filled kernels | Appearance of brown rice |  |  | Kernel dry weight |  |  |  |
| --- | --- | --- | --- | --- | --- | --- | --- | --- |
|  |  | Perfect kernels | White-back kernels | Other kernels | Perfect kernels | White-back kernels | Other kernels | Average |
|  |  | % |  |  | *mg · kernel<sup>-1</sup>* |  |  |  |
| 26°C | 97.9 | 100.0a | 0.0b | 0.0a | 21.6 | - | - | 21.7a |
| N+26°C | 96.5 | 98.9a | 0.0b | 1.1a | 21.8 | - | 22.7a | 21.8a |
| 34°C | 94.4 | 11.4c | 66.4a | 22.2a | 19.9 | 20.2 | 20.2b | 20.2b |
| N+34°C | 94.7 | 52.1b | 7.6b | 40.4b | 20.2 | 20.7 | 20.0b | 20.2b |

Values in appearance are means measured from superior kernels attached to the upper position of a panicle from 3-5 panicles in each treatment. The dry weight of each kernel type was means of individually collected 3-7 panicles, and the average dry weight was calculated in each treatment. Means within a column followed by different letters indicate a significant difference ( $p < 0.05$ ) (Tukey's Honestly Significant Difference [HSD]).

**Figure 1.**
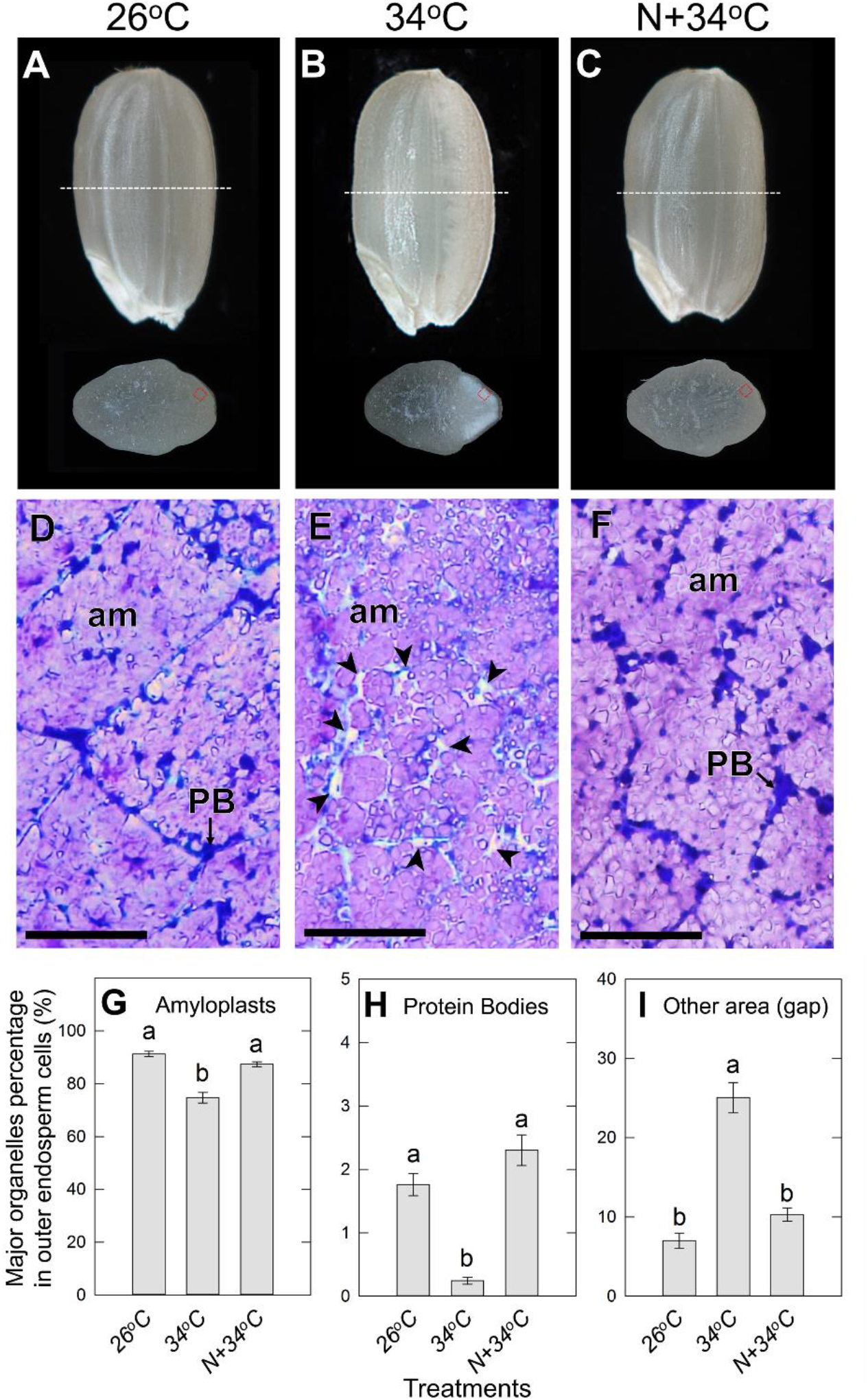
Images of rice kernel and the transverse sections of 26°C (A), 34°C (B), and N+34°C (C) treatments at maturation (40 DAH). Light microscopic images taken in the vicinity of the outer endosperm in the dorsal side of kernels harvested in 26°C (D), 34°C (E), and N+34°C (F) treatments, corresponding to the red dotted square on the transverse sections inserted in A-C. Grain sections of D-F were double-stained with Coomassie brilliant blue and iodine-potassium iodide. In D-E, PB, protein body; am, amyloplast. Arrows in E indicate gap areas. Bars in D-F indicate 50 μm. Area-based percentages of major organelles, amyloplasts (G), protein bodies (H), and other area (gap) (I) occupying in the outer endosperm cells in 26°C, 34°C, and N+34°C treatments were shown. The data indicate that the means±SE of 11-25 individual cells collected from at least 3 kernels from 3 individual plants in each treatment. Different letters in G-I indicate a significant difference (Tukey’s HSD test, *p* < 0.05).

### Microscopic Observations in Chalky Zone

The outer endosperm cells located at the dorsal side, where the high frequency of chalkiness was confirmed at maturation (Table1), were observed microscopically using a light microscope and TEM. Rice grain sections collected in 26°C, 34°C, and N+34°C treatments were dual-stained and observed using a light microscopy, and starch granules and proteins were stained in bluish purple and blue area, respectively (Figures 1D-F). In 26°C treatment, the outer endosperm cells were densely packed mainly with numerous well-developed amyloplasts and mature PBs mostly located at the periphery of cytoplasmic compartments (Figure 1D). Heterogeneity in the size of amyloplasts was contrastingly observed in 34°C treatment as well as a relatively large gap space in the cytoplasmic compartments (see the arrowheads in Figure 1E). Cell morphology in N+34°C treatment was similar to that in 26°C treatment (Figure 1F). The image analysis showed that 34°C treatment significantly reduced areas of amyloplasts and PBs in the cytoplasmic compartments (Figures 1G and H) and resulted in the preservation of over 25% of gap spaces among amyloplasts and PBs inadequately formed in the chalky zone, different from the perfect kernels in 26°C treatment (Figure 1I). In N+34°C treatment, the area-based percentage of amyloplasts and PBs recovered at the similar level to the 26°C treatment (Figure 1G-I). Figure 1I indicates that these modifications of organelle compartmentations observed at high temperature were ameliorated by additional N supply (Figure 1F) and displayed no difference in 26°C treatment (Figure 1I).

In the outer endosperms in 34°C treatment, the gap area was typically located among amyloplasts in the cytosol. Microscopic observation was then made by using TEM in the same zone for each treatment at 12, 20, and 40 DAH (Figures 2A-I). At 12 DAH, the developing PSVs were similarly observed in all treatments. Numerous PSVs, few PBs, mitochondria, and rERs were observed in the cytosol (Figures 2A-C). In contrast to 12 DAH, the ultrastructure of the cells fixed at 20 and 40 DAH both differed between treatments. The majority of PSVs were filled with PBII until 20 DAH in 26°C treatment (Figure 2D), whereas PSVs in 34°C treatment remained in the cytosol, leading to the formation of air spaces at maturation (Figures 2E and H). In N+34°C treatment, normal PBII development was observed with little or no PSVs at 40 DAH, similar to 26°C treatment (Figure 2I). In each treatment, the water content of the kernels decreased through the grain-filling stage, although at maturation, higher water content was observed in 34°C treatment, compared to other two treatments (Figures 2J-L).

**Figure 2.**
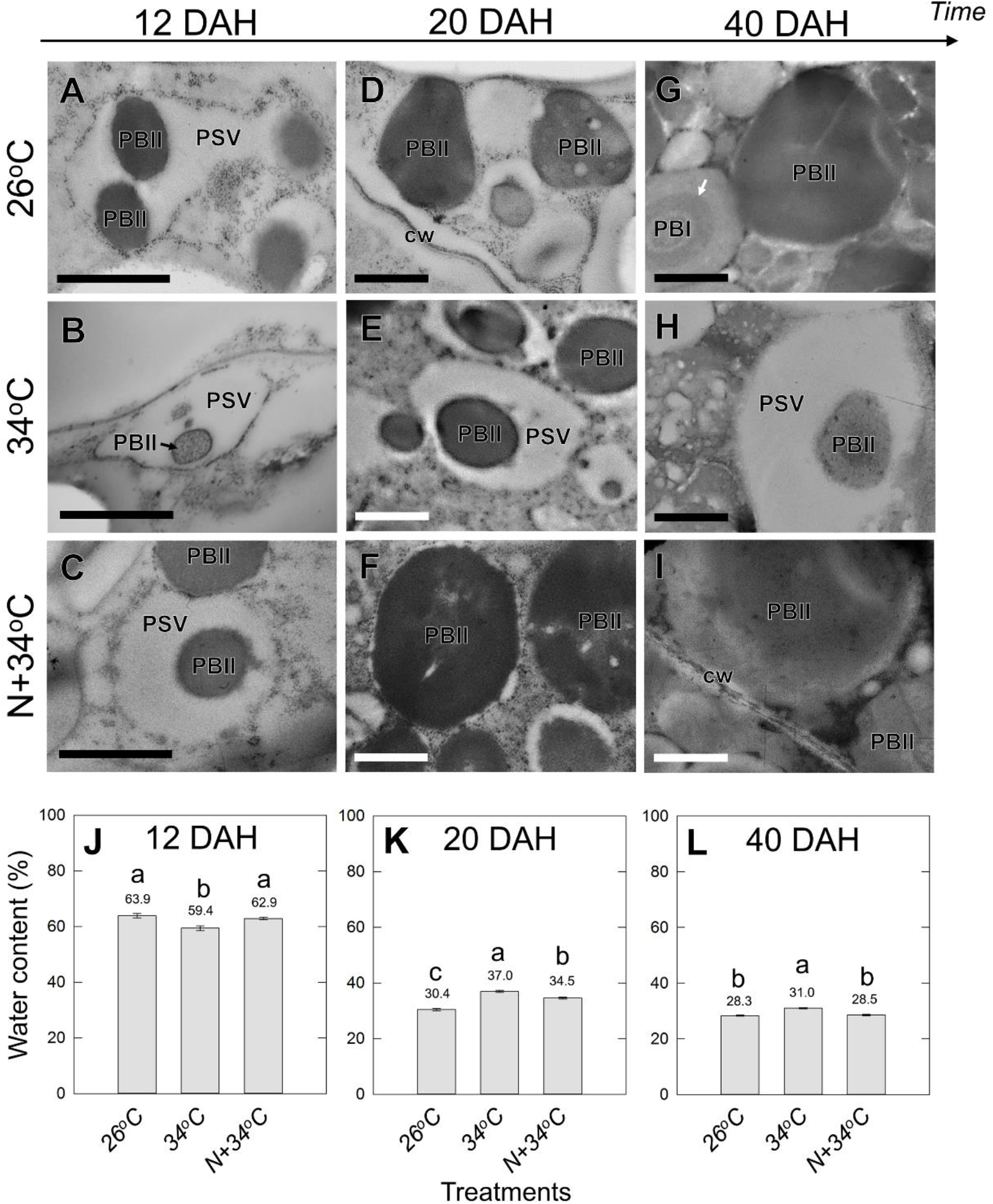
TEM images of the outer endosperm cells in 26°C (A, D, G), 34°C (B, E, H), and N+34°C (C, F, I) treatments collected at 12 (A-C), 20 (D-F), and 40 DAH (G-H). Water contents of rice grain kernels in each treatment at same three ripening stages are shown (J-L). Data are means±SEs of 18 kernels collected from 3 plants, and means are shown on the bars in each figure. Different letters indicate a significant difference among treatments (Tukey’s HSD test, *p* < 0.05). In A-I, PSV, protein storage vacuole; PBI, protein body type I; PBII, protein body type II; cw, cell wall; rER, rough-surfaced endoplasmic reticulum. In G, each arrow indicates the deep-colored areas of Cys-rich 10-kDa layer formed in PBI. Bars indicate 1 μm.

### *In Situ* Turgor Assay in Outer Endosperm Cell Growing under Heat Conditions

In the corresponding target cells at the early stage, where PSVs were similarly localized in all treatments (Figures 2A-C), there were some kernel-to-kernel variations in outer endosperm cell turgor in each treatment (Figure 3B) because each kernel exhibited some variations in the appearance at maturation (Table 1). 34°C-treated cells exhibited positive but relatively low turgor, compared to those in 26°C treatment (Figure 3B). Cell turgor in N+34°C treatment was overall higher in 34°C treatment, maintaining the same level that of 26°C treatment (Figure 3B).

**Figure 3.**
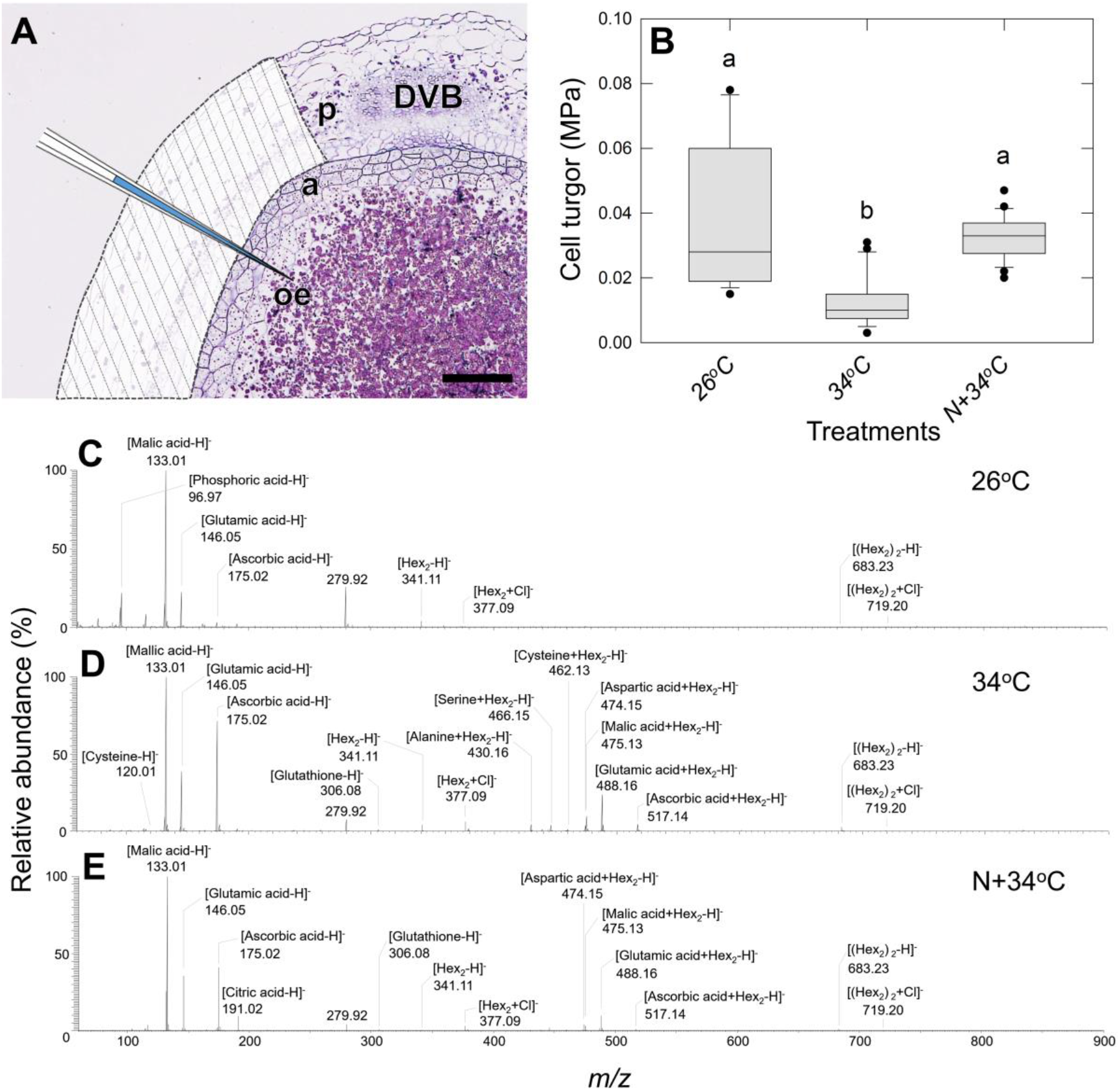
Image of the transverse section of the dorsal side of rice kernel at the early ripening stage (12 DAH) (A). In A, an image of cell sap extraction using a pressure probe from the outer endosperm cells was also drawn. A part of the pericarp shown in white with oblique lines was removed prior to the insertion of a probe tip, and a cell pressure probe was introduced (see Materials and Methods). DVB, dorsal vascular bundle; a, aleurone layer; oe, outer endosperm; p, pericarp. Bar indicates 100 μm. In B, box plot illustrating the variations of outer endosperm cell turgor in each treatment assayed at the same stage. Data are mean±SEs for 14-29 cells from 7-8 kernels from at least three different plants in each treatment. Different letters indicate a significant difference among treatments (Tukey’s HSD test, *p* < 0.05). C-E, PicoPPESI-MS negative ion mode mass spectra obtained from the cells in 26°C (C), 34°C (D), and N+34°C (E) treatments at the same stage. Data are representative of similar experiments with 4-5 kernels in each treatment.

### On-site Cell-specific Metabolites Profiling in Kernels Growing under Heat Conditions

When the cell sap was directly analyzed using picoPPESI-MS under controlled environments (Supplementary file 1), numerous metabolites (mostly amino acids, sugars, organic acids, and secondary metabolites) were identified in negative ion mode at less than 5 ppm differences from theoretical molecular weight values (Figures 3C-E, Supplementary file 2). In 26°C treatment, the peaks of phosphoric, malic, and glutamic acids (as [M−H]^−^, M=molecular species), and some sugars (as [M−H]^−^ and/ or [M+Cl]^−^) were observed in the mass spectrum as major ions (Figure 3C, Supplementary file 2). Under 34°C treatment, the signal intensity of glutamic acid was much higher than in 26°C treatment, and the content of ascorbic acid, glutathione, and amino acids increased as in typical heat responses (Figure 3D, Supplementary file 2). The signal intensities of ascorbic acid and glutathione both involved in the scavenging of reactive oxygen species (ROS) were higher in 34°C treatment than in 26°C treatment (Figure 3D; Supplementary file 2). Moreover, the accumulation of cysteine (Cys) that forms the disulfide bridges was observed with over 75% of frequency under 34°C treatment, whereas Cys-related signals were not found in 26°C treatment (Supplementary file 2). In N+34°C treatment, the signal of Cys as [Cys−H]^−^ was rarely observed (<20%), consistently with those of Cys-sugar cluster ions, such as [Cys+Hex−H]^−^ and [Cys+Hex_2_−H]^−^, and generated during the ionization process (not detected and 0.09%, respectively; see Supplementary file 2). Higher sugar accumulation than heat was observed under N+34°C treatment (Figure 3E). For methionine (Met) which forms disulfide bonds similarly to Cys, the signal intensities of [Met−H]^−^ and two Met-sugar cluster ions, [Met+Hex−H]^−^ and [Met+Hex_2_−H]^−^, were smaller than those of Cys, with no clear differences in the different treatments (Supplementary file 2).

### Heat and Nitrogen-enhanced Adaptation on Protein Body Development in the Endosperm Cells

When exposed to heat, the weight of proteins per kernel decreased. However, the protein weight per kernel in N+34°C treatment was similar to that in 26°C treatment (Figure 4B). The number and area of each PB decreased remarkably in 34°C treatment (Figures 4C-F), and N application increased the number of each PB and area of PBII, but not completely for the area of PBI (Figures 4C-F). The area of PBII in 26°C treatment was approximately 2.5-fold greater than PBI localized in the cells (Figures 4E and F). PBI is composed of two types of layers, namely, the inner Cys-rich 10-kDa prolamins (CysR10P) layer and outer layer containing the mixture of other prolamins (see Figures 2G and I). In 34°C treatment, the apparent area of CysR10P in PBI increased, although the outer layer was decreased remarkably (see deep-colored bars in Figure 4E). In N+34°C treatment, the area of CysR10P in PBI was similar to that in 34°C and prone to be higher than 26°C treatment. Consequently, the area of CysR10P (number x area) in the chalky cells increased due to N application, compared to that in 34°C treatment (Supplementary file 3). There was little effect on the accumulation of glutelin precursors (pro-glutelin) and acidic (α)-glutelin to 34°C, although the content of CysR16P, Cys-poor 13-kDa prolamin (CysP13P), α-globulin, and basic (β)-glutelin decreased in 34°C treatment (Figure 4G). When N was supplied to the soil prior to 34°C treatment, the content of β-glutelin and CysR16P increased to the level of 26°C treatment and greater than that of 34°C treatment (Figure 4G). The accumulation of CysR10P was specifically observed in 34°C treatments (Figure 4G). Analysis of the time course of changes in PSV and PBII (stored in PSV) volumes in the chalky zone showed that the expansion of PSVs in 34°C treatment progressively occurred, reaching 26 fL at maturation, 2.6-fold larger than that in 26°C treatment (Figure 5A). PB accumulation rate retarded in 34°C treatment, compared to that in 26°C treatment (Figure 5B). Additionally, in 26°C and N+34°C treatments there were positive and linear relationships between the entire PSV and PB volumes (inset in Figure 5C). In contrast, the 34°C-treated PSV volume was shown to be positively correlated with the PSV matrix volume (Figure 5C).

**Figure 4.**
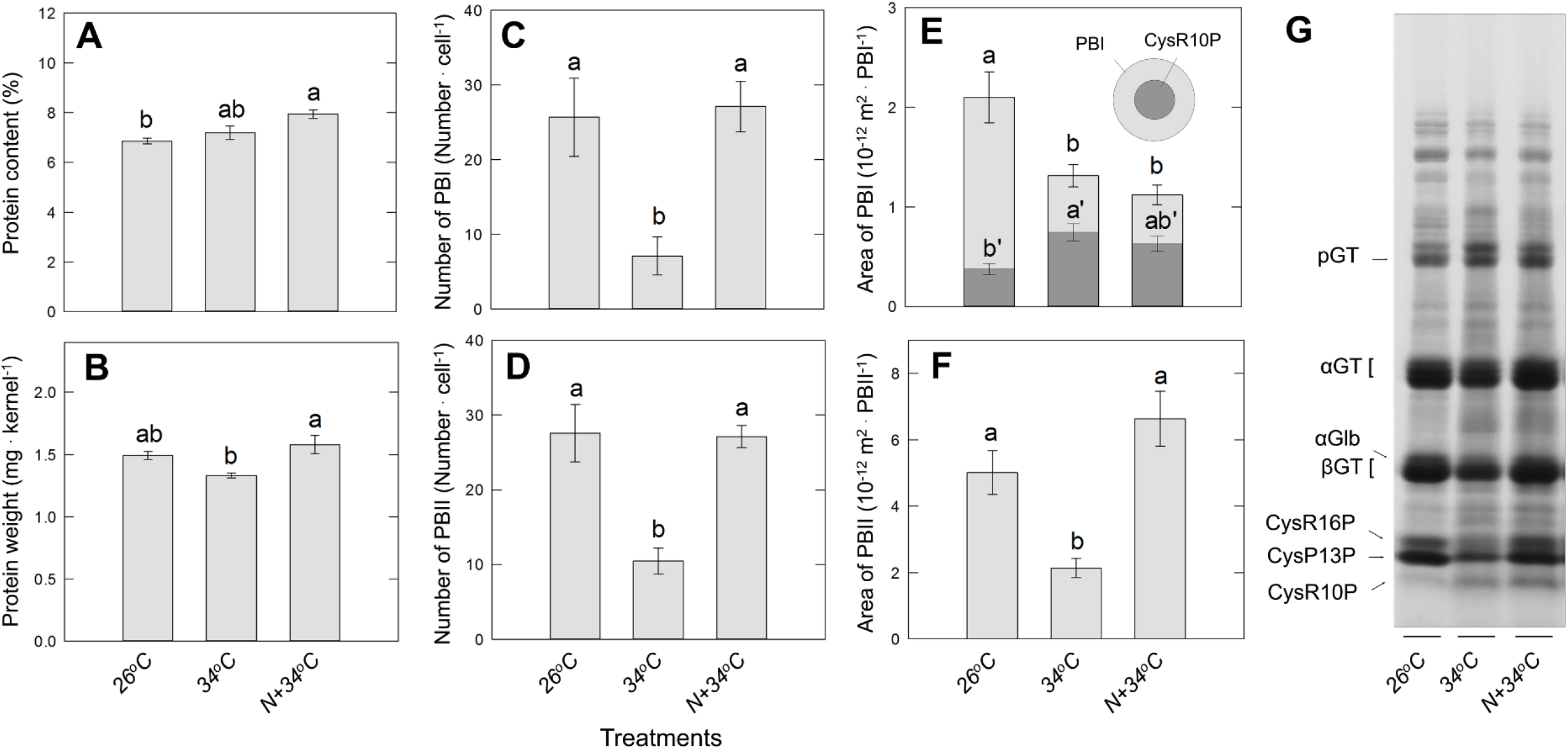
Protein content (A) and protein weight (B) of kernels in each treatment at 40 DAH. Data in A and B are mean±SEs for four plants. C-F, the number (C, D) and cross section area (E, F) of PBI (C, E) and PBII (D, F) in the outer endosperm cells morphometrically analyzed with the TEM images at 40 DAH (see Figure 2G, H, I). Each parameter in C-F indicates mean±SEs for 25-59 PBs from at least three kernels. In E, each deep-gray bar embedded in the light gray bars showing the area of PBI indicates the area of Cys-rich 10-kDa layer, which locates at the core of PBI (see inset). Different letters in A-F represents significant differences among treatments (Tukey’s HSD test, *p* < 0.05) within the protein content, protein weight, and the area and number of both PBs (letter only) and area of Cys-10-kDa layer of PBI (’). In G, SDS-PAGE analysis of one-third of dorsal side of kernel, corresponding to the chalky zone in 34°C treatment. CysR10P, Cys-rich 10-kDa prolamin; CysR16P, Cys-rich 16-kDa prolamin; CysP13P, Cys-poor 13-kDa prolamin; pGT, proglutelins; αGT, glutelin acidic subunit; βGT, glutelin basic subunit; αGlb, α-globulin.

**Figure 5.**
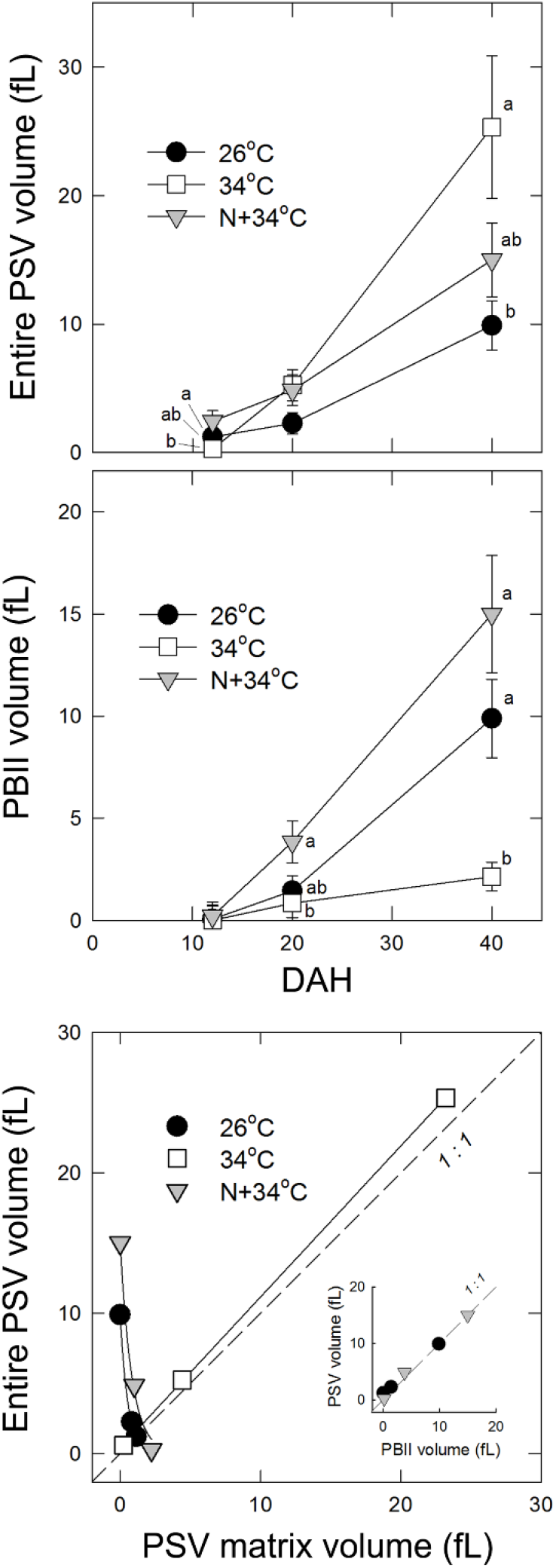
Time course of changes in the estimated PSV (A) and PBII (B) volumes in each treatment. In C, the PSV volume as a function of the corresponding PSV matrix volume. Closed circle, opened square, and gray downward-pointing triangle indicate 26°C, 34°C, and N+34°C treatments, respectively. Each point indicates mean±SEs calculated from areas of 4-38 PSVs or PBIIs. In A and B, nitrogen application at 4 DAH was shown, and the thick black bars at the x-axis indicate the duration of 34°C treatment. Different letters in A and B indicate significant differences among treatments (Tukey’s HSD test, *p* <0.05). In C, the regression line between the PSV matrix volume (*x*) and the entire PSV volume (*y*) in 34°C treatment was *y* = 1.08*x* +0.41 with *r*^2^ = 0.99 (*p* < 0.01). Inset in C shows the relationship between PSV volume (*y*) and PBII volume (*x*) in 26°C and N+34°C treatments, and the regression lines in 26°C and N+34°C were y=0.89x with *r*^2^=0.99 (*p* < 0.01) and y=0.98x+0.01 with *r*^2^=0.99 (*p*<0.05), respectively. Each dashed line in C indicate a 1:1 line.

## Discussion

Rice chalkiness is a typical physiological disorder of immature kernels and it exhibits air spaces between loosely-packed starch granules causing a random light reflection (Tashiro and Wardlaw, 1991b; Zakaria et al., 2002). Deterioration in the milling quality due to an increase in heat-induced chalky rice has been a serious concern in global rice production under the advancing global warming. N supply decreases heat-induced chalky formation (Perez et al., 1996; Wakamatsu et al., 2008), although the underlying mechanisms of heat-induced chalky formation and the N-enhanced adaptation have never been systematically examined mostly due to the technical difficulties at the cellular level. As pointed previously (Hakata et al., 2017), another difficulty for studying heat-related damages in crop plants would be due to the low reproducibility of high-temperature treatment under field conditions, which implies the need of environmental controls. We have developed a new on-site cell metabolomics performable in the controlled environments to make such a cellular analysis possible (Supplementary file 1). By using this method, we carried out a cell-specific analysis in the putative chalky zone in the attached kernels grown under high-temperature environments, in a combination with a transmission electron microscopy. It has been uncovered that the preservation of relatively large PSVs among loosely-packed starch granules in the cytosol was attributed to the air spaces formed in the dorsal chalky zone in the kernels. Although 34°C-treated cells disturbed protein synthesis, N supply enhanced the protein synthesis rate to sustain normal development of amyloplast and PBs in the zone, leading to the reduction in air spaces under heat conditions. Therefore, we conclude that rice endosperm cells could alter vacuolar morphology by regulating vacuolar trafficking and protein processing as a heat stress adaptation, depending on the available nitrogen level. We also propose that rice chalky formation is a form of heat acclimation.

### Establishing an on-site cell-specific analysis to investigate heat-induced damages in rice plants

Recently, single cell metabolomics has been extended mainly due to the introduction of Orbitrap mass analyzer, and this approach has been applied to a number of biological studies including plant cell analysis (Gholipour et al., 2013; Fujii et al., 2015). A cell pressure probe, originally invented by Steudle’s group (Hüsken et al., 1978) and long-used to measure the cellular water status in plant sciences, has been adopted as a picolitre pipette to establish a new type of *in-situ* analytical method by combining with an Orbitrap mass spectrometer in Nonami’s laboratory (Gholipour et al., 2013). More recently, both resolution and sensitivity of the analytical method have been improved by introducing an internal electrode in the capillary holder and adopting a mixture of an ionic solution and silicone oil in the quartz capillary (Nakashima et al., 2016). This method, called ‘picolitre pressure-probe-electrospray-ionization mass spectrometry (picoPPESI-MS)’, is performable in intact plants. This type of cell metabolomics appears to be a powerful tool. However, to the best of our knowledge, most existing cell metabolomics, including this method, has been confined to the laboratory use to study cellular metabolisms at room temperature. To date, no attempts have been made on further improvement of the methods to investigate metabolic responses to some environmental stimuli, such as heat responses in developing crop plants that we examined here.

To meet the requirement, a new on-site cell-specific analytical method has been developed by combining picoPPESI-MS with environment controls (Supplementary file 1). In this system, both developing plants and analytical environment could be equilibrated at the same set temperature. Under temperature equilibrium conditions, a probe tip was finely introduced into the target endosperm cells prior to chalky formation. The picolitre cellular fluids discharged into the quartz capillary were collected, and the metabolome analysis was carried out directly without any pretreatment (see Materials and Methods section). The use of the analytical method at the early ripening stage allowed determination of the differences in metabolites between treatments in the developing dorsal outer endosperm cells, where numerous PBs are known to be localized (Ellis et al., 1987; Hoshikawa, 1989) (Figure 3). As discussed below, it is noteworthy that our cell-specific analysis clarified that the fate of the endosperm cells was determined, in which there were no obvious morphological treatment differences (Figure 2A-C).

### Direct evidence for morphological and metabolic characteristics of chalky formation and nitrogen-enhanced adaptive process under heat conditions

Figure 6 shows the likely processes of chalky formation and the N-enhanced adaptation occurring in the outer endosperms cells under heat conditions. Our results provide a new evidence that at high temperature, the volume of PSVs increased through maturation and remained in the cells at maturation, with a limited accumulation of storage proteins, mostly glutelin and globulin (Figures 4 and 5). When N was supplied prior to 34°C treatment, the synthesis of proteins including those storage proteins was sustained under heat conditions, resulting in normal PBII development (Figures 2 and 4E), accompanied with a substantial starch accumulation (Figures 1G and 6) and PBI development (Figures 4C and E, Supplementary file 3). Under N+34°C treatment, the spatial ratio of the gap spaces at maturation was similar to that of 26°C treatment, 14.8% lower than that of white-back rice harvested in 34°C treatment (Figure 1I). As the results indicated, approximately 70% (Δperfect rice 40.7% · Δwhite-back rice 58.8%^−1^) of white-back rice simply turned into the perfect rice without causing any reduction in kernel weight (Table 1).

**Figure 6.**
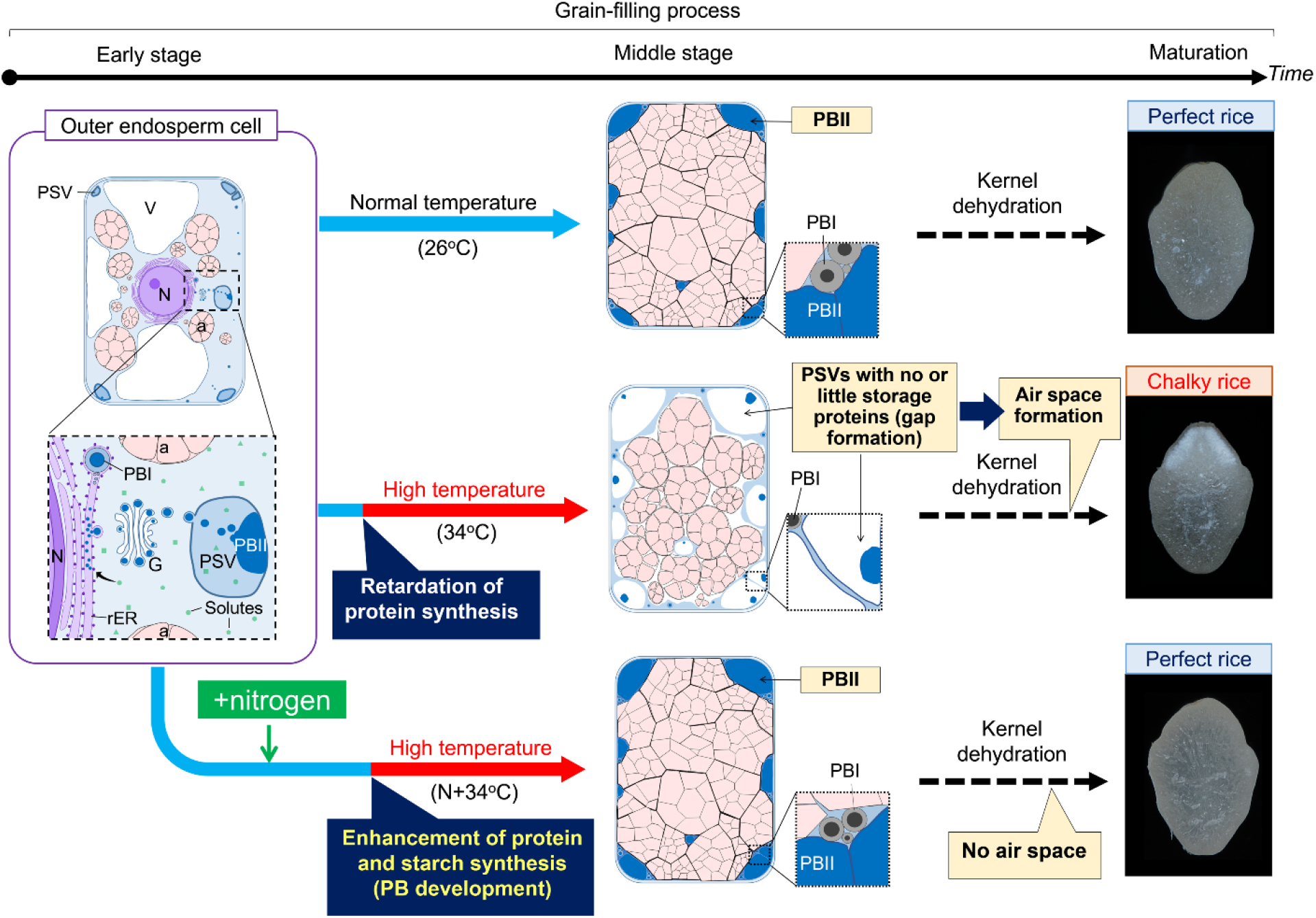
Schematic diagrams of the putative process of heat-induced chalky formation and nitrogen-enhanced adaptation in the dorsal outer endosperm cells. In the cells under normal conditions (26°C treatment), as the packing density of amyloplasts in the cells increases according to the reduction in the cytosol occupied with mainly vacuoles, resulting in the adequate development of amyloplasts and PBs to diminish cytosolic space (see Supplementary file 4C, F and I). When N level is low prior to high temperature (34°C treatment), the cells adjust less osmotically and reduce protein synthesis rate to preserve PSVs and vacuoles (see Discussion), resulting in air space formation during kernel dehydration, which turns to chalk. When N supply is adequate in the cells prior to high temperature (N+34°C treatment), cells sustain protein synthesis rate by strong osmotic adjustment to promote normal PB and amyloplast development, which diminishes cytosolic space, resulting in a substantial increase in perfect rice.

It has been shown that heat stress affected the PB development, and the volume of PSVs progressively increased over time, reaching to 2.5-fold larger than PBII in 26°C treatment at maturation (Figures 4F and 5A and B). Based on the observed size and number of each PB (Figures 4C-F), our calculation suggests that the volumetric ratio of PSV to cytosolic volume corresponded to 24.5% on average (ranging between 6.5–46.4%) of cytosol, indicating that PSVs would be a predominant compartment occupying the cytosol at least at maturation. Hence, the observed spatial modifications of vacuolar compartmentation (Figures 1E and 6) account for the gap space formation under heat conditions. Most mutants with abnormal PBII formation showed floury endosperms, whose appearance was opaque (Takemoto et al., 2002; Wang et al., 2010; Fukuda et al., 2011; Ren et al., 2014), whereas a vacuolar processing enzyme, *OsVPE1* knockdown mutant contrastingly produced translucent kernels (Wang et al., 2009). Given the fact that OsVPE1 plays a role in the maturation of glutelin transported into PSV as Cys protease in the final stage of PB development (Wang et al., 2009), it is not surprising that unlike other enzymes, VPE does not have a dramatic volumetric impact on chalky formation. Thus, the results of Wang et al. (2009) do not necessarily conflict with our conclusion.

Osmotic adjustment occurs in the growing cells at moderately low water potential (Morgan, 1977; Cutler et al., 1980a, 1980b; Meyer and Boyer, 1981). During the osmotic adjustment, turgor pressure can be maintained by accumulating osmotically active solutes, such as sugars and amino acids, into the cells (Morgan, 1977; Meyer and Boyer, 1981). As reported previously (Wada et al., 2011; Wada et al., 2014), osmotic adjustment also occurred in the rice endosperms cells growing under dry wind conditions prior to chalky ring formation. On-site cell metabolomics and *in situ* turgor assay conducted in outer endosperms indicated clear treatment differences in the heat adaptive responses (Figure 3B-E). Different from the drought-related studies, there seem to be few studies regarding osmotic adjustment under heat conditions (e.g., Jiang and Huang, 2000 in turfgrass). In this study, the determination of cell osmotic pressure was not attainable using a freezing point osmometer, because of the interference of starches contained in the sap. However, based on the treatment differences in metabolites in the target zone, it was reasonably interpreted that N-treated cells strongly adjusted osmotically to maintain the protein synthesis rate at high temperature, whereas under low N level, the cells adjusted less osmotically and the protein synthesis rate slowed down, and therefore the energy requirement in cytosol could be maintained low.

Cys stabilizes the tertiary structure of proteins through the formation of disulfide bonds, which is necessary for protein folding and the modulation of enzyme activity in the cells. N-induced increase in Cys-rich 10-kDa prolamin in PBI (Figure 4E) and the result of SDS-PAGE (Figure 4G) are strong evidence supporting our conclusion. The results of cellular metabolome analysis suggest that during the N-enhanced adaptation process, Cys would have contributed to vacuolar trafficking and protein processing without participating in osmotic adjustment as an osmotically active solute. The greater abundance of Cys-related signals was frequently observed in the heat-treated cells, but no or less in other two treatments (see [Cys−H]^−^, [Cys+Hex−H]^−^, and [Cys+Hex_2_−H]^−^ in Supplementary file 2). And, this data corresponds to the occurrence of white-back kernels (Table 1). Hence, it is expected that higher Cys accumulation would have occurred in the cytosol in 34°C treatment, as well as sugars and other major amino acids, which might be caused by a low assimilate input under long-term heat conditions. It appeared that N application increased amyloplast development in the cells under heat conditions (Figure 1G), and hence the effects of N supply were large enough to structurally stabilize the activity of a series of starch biosynthesis-related enzymes under heat conditions, together with PB development.

For nitrogen budget in the cells, it remains questionable how much N had been synthesized into the storage proteins during the adaptation process. In the comparative analysis between 34°C and N+34°C treatments, our TEM image analysis indicated that the treatment difference in the spatial ratio of PBI and PBII occupied in the cells was 0.52% and 3.86% on average, respectively. The percentage of the chalky area to the transversal area corresponds to 17.4 ± 0.8% (mean ± SD, *n* = 18). The mean area of the transversal section measured in 34°C- and N+34°C-treated kernels was 0.051 and 0.050 cm^2^, respectively, and the averaged longitudinal kernel length in these treatments was 5.15 and 5.16 mm, respectively. With the specific gravity of PBI and PBII, 1.27 and 1.29 g·mL^−1^, respectively (Tanaka et al., 1980), the weight difference in the chalky zone between those treatments corresponded to 0.32 mg·chalky zone^−1^. Because the treatment difference in the protein content was 0.25 mg·kernel^−1^ (Figure 4B) and the difference in the occurrence of white-back rice was 58.8% (Table 1), our estimation suggests that in N+34°C treatment approximately 75.1% of proteins increased by N application were synthesized as storage proteins in the PBs to fill up the gap spaces in the chalky zone. However, the rest of 24.9% was consumed for synthetizing PBs in other area and other enzymes, mostly for starch synthesis.

### The adaptation strategies of rice endosperms to heat conditions

There is a large body of evidence that ROS, such as hydrogen peroxide, play a role as important second messengers in signal transduction networks associated with developmental processes or in response to abiotic stress (Mittler et al., 2004; Skopelitis et al., 2006). Generated hydrogen peroxide might serve as a signal introducing the programmed cell death and subsequently kernel desiccation (Onda et al., 2009). More recently, Ibl et al. (2014) suggested that both the membrane shrinkage and degeneration of PB occurred before PB development were completed in barley endosperm cells. The synthesis of vast amounts of disulfide-rich storage proteins during grain-filling (Figure 4E and G) might be accompanied by the production of hydrogen peroxide in ER, resulting in the peroxidation of membrane lipids under normal conditions (Sattler et al., 2006; Onda et al., 2009). In contrast, the behavior of PSVs we observed (Figures 2H and 5) suggests that heat might have disturbed the cellular redox status to inhibit the peroxidation of membrane lipids under heat conditions. PSVs treated at 34°C were found to be expanding over time, but with a retardation of PB accumulation (Figure 5A and B). Importantly, the volumetric increase in PSV could be explained by increasing PSV matrix (see Figure 5C), indicating that substantial amount of water had been entering PSVs towards maturation to increase the vacuolar volume. Thus, it is quite unlikely that tonoplast membrane lipids were degraded under heat conditions. One plausible explanation is that a partial degradation of PBII in PSV would occur through the activation of proteases or the autophagy-like process, leading to an increase in the vacuolar osmotic pressure. This would promote water entry into PSVs to sustain the vacuolar volume, and maintain the cell volume. The source of water accumulated in the PSV matrix remains unknown. However, based on the increases in the content of ascorbate, glutathione, and monodehydroascorbate detected at the cellular level (Supplementary file 2), we speculate that the accumulation might be a consequence of an increased activity of ascorbate peroxidase catalyzing the conversion of hydrogen peroxide into water. PB development sustained during the N-enhanced adaptation process supports our conclusion that disulfide bond formation and tonoplast denaturation would be facilitated by strong osmotic adjustment (Figure 2). As observed in Figure 2J-L, 34°C-treated kernels exhibited higher water content than 26°C treatment, consistent with previous studies (Ishimaru et al., 2009; Iwasawa et al., 2013; Hayashi et al., 2015). Given the fact that water is a major compound in both PSV matrix and vacuoles in the gap spaces, it is not surprising that chalky zone (or even chalky rice) exhibited relatively high moisture content under heat conditions, compared to 26°C treatment. Storing water in the endosperms along the dorsal vasculature may be an essential event to sustain embryo development in the seeds exposed to the extremely high temperature, as heat-induced precocious germinations were similarly observed in rice (Iwasawa et al., 2013) and oilseed rape seeds (Brunel-Muguet et al., 2015). Hence, we propose that chalkiness is a form of acclimation to heat stress.

### Threshold of chalkiness and other possible candidate organelle involved in chalkiness

Regarding the threshold above which chalkiness appears, our image analysis suggests that the transition between chalk and translucence corresponds to a range between 10.3 and 25.1% in N+34°C treatment and 34°C treatment, respectively (see Figure 1I). In another type of chalkiness, called milky-white rice, chalkiness was observed at 13.3% on average (Wada et al., 2014). Taken together, significant transparency loss would occur in a range of 10.3 to 13.3%. The preservation of PSVs expanded in cytosol was responsible for the formation of heat-induced white-back rice (Figure 6). In the chalky zone, a small number of cells, corresponding to 3.4% of the total chalky area (see the red area in Supplementary file 4A) and located adjacent to the sub-aleurone layer, retained some large lytic vacuoles in addition to PSVs (Supplementary file 4H). This observation has risen a possibility that in ‘Koshihikari’, some vacuoles might also have participated in the air space formation. Similarly, we previously showed that the presence of vacuoles stayed in the cytosol at osmotic adjustment might have an impact on the formation of air spaces in the inner endosperm cells in a foehn-induced chalky ring formation (Wada et al., 2014). Wakamatsu et al. (2008) reported a clear varietal difference in the occurrence of white-back rice when the same amount of N was applied under heat conditions; however, the underlying mechanisms remain unclear. Some morphological differences may explain the difference that is currently under investigation. More recently, it has been possible to trace metabolites in the target zone using the stable isotope(s) in mass spectrometry (Wada et al., 2017). The use of this analytical method may extend our understanding of heat adaptation mechanisms in rice endosperms.

### Conclusion and remarks

In this work, the on-site cell-specific analysis was utilized to investigate the cellular mechanisms of heat-induced chalky formation and N-enhanced adaptation response. Combined analysis with a TEM demonstrated that the cells adjusted less osmotically and reduced protein synthesis to retain PSVs in the cells at low N level under heat conditions, and that N supply enhanced the heat tolerance and the cells sustained protein synthesis during strong osmotic adjustment at high temperature. During the N-enhanced adaptation process, the formation of disulfide bonds was promoted by consuming Cys to sustain normal PB and amyloplast development even under heat conditions, which resulted in a remarkable increase in perfect rice. Hence, the extent of translucent loss in endosperms could result from the spatial modifications of organelle compartments, closely associated with available N level in the cells. Our on-site cell-specific analysis can be applied in a number of cell-specific studies in future, including heat-induced rice spikelet sterility. Finally, to our knowledge, no attempts have been made in other instruments such as a nuclear magnetic resonance. However, a similar combination with environmental controls may be useful in the field of environmental plant biology, particularly when investigating temperature stress responses in crop plants, as reported here.

## Materials and Methods

### Plant materials

*Oryza sativa* L. ‘Koshihikari’ potted plants were grown without giving a top dressing in Kyushu Okinawa Agricultural Research Center, Chikugo, Japan, according to the method of Wada et al. (2011). At flowering, the potted plants grown in field were transferred to the walk-in environmentally-controlled growth chamber (26/22°C, 70/80% relative humidity, and 750 μmol photons m^−2^ s^−1^ photosynthetically active radiation [PAR]) set at the plant canopy using light emitting plasma (LEP) lamps (STA 41-02, Stray Light Optical Technologies, Inc., IN, the US) with a photoperiod of 13 h day/11 h night. For half of the potted plants, 0.45 g·pot^−1^ of urea was applied at 4 DAH to examine the effects of N application under heat conditions on rice quality. At 5 DAH, half of the plants were transferred to another growth chamber set at 34°C and 70% relative humidity (RH) (09:00-15:00) / 28°C and 80% RH (15:00-09:00) and 750 μmol m^−2^ s^−1^ PAR with the same photoperiod to be treated at high temperature for 10 days (referred to as ‘34°C treatment’). Other potted plants were kept in the same chamber in a non-heat (26°C) treatment. At 15 DAH, plants were transferred to the control chamber to grow until the mature stage (typically 40 DAH).

### On-site cell metabolomics and turgor assay

On-site live cell metabolome analysis using picolitre pressure-probe-electrospray-ionization mass spectrometry (picoPPESI-MS, Nakashima et al., 2016) was carried out in the dorsal outer endosperm cells of the superior kernels attached to the upper position of a panicle, where a high frequency of chalkiness was observed under 34°C treatment (Table 1). Simply, the system was composed of picoPPESI-MS and two measurement rooms individually attached to growth chambers (K260B029-S01, Tsubuku Corporation Ltd., Kurume, Japan) (see Supplementary file 1). The combination of the environmental control and picoPPESI-MS allowed us to directly perform cell metabolites profiling real-time at the target zone without any pretreatments under each set environmental conditions. Potted plants at 11–12 DAH were moved to the measurement room where they were placed at the center of a U-shaped vibration-free table (HOA-0808LA(Y), Herts Co. Ltd., Yokohama, Japan) in the measurement room. Temperature change in the growth chamber disturbs cell pressure probe measurement (Boyer, 1995), and therefore the following assays were initiated, at which the pressure probe system and plants reached to the temperature equilibrium (typically in 30 min). Prior to the assay, a part of the hull (i.e., palea) in the attached kernels was quickly and gently removed under humid conditions (Wada et al., 2011). In the preliminary experiment, it was observed that there were distinct tissue-to-tissue variations in metabolites between the pericarp (Supplementary file 5) and outer endosperm cells (Figures 3C-E). When the kernel score reached to 0.9 (see Figure 1 in Wada et al., 2014), a 1 mm diameter biopsy punch was used to remove 0.031 cm^2^ of pericarp tissue in the dorsal side of the kernel to obtain the sap from the endosperm cells under humid conditions prior to the capillary tip insertion (Figure 3A). Therefore, a possible contamination could be ruled out from the pericarp cell layers. The kernel was gently fixed on the sample holder using tape and magnets (Supplementary file 1). The tip of microcapillary filled with 0.01% (v/v) ionic liquid/silicone oil mixture was impaled into endosperm cells (typically between 50-150 μm below nucellar-epidermis) with the aid of a motorized piezomanipulator (DC-3K, Märzhäuser Wetzlar, Germany). Cell sap was collected by depressurizing in the microcapillary, and the tip immediately oriented toward the orifice of an Orbitrap mass spectrometer (Q-Exactive, Thermo Fisher Scientific Inc., MA, the US) was electrified with −4 kV using a high voltage generator (AKTB-05k1PN/S, Touwa Keisoku Corp., Tokyo, Japan). The MS scan was performed in negative ion mode with the instrumental settings of 200 ms as maximum injection time, inlet ion transfer tube temperature of 250°C, and resolution of 35,000. When the target cells were successfully impaled to collect picolitre cell sap without tip plugging, the entire process of picoPPESI-MS analysis on the cells was completed within few minutes. All the standard chemicals and organic solvents used in the experiments were liquid chromatography-mass spectrometry (LC/MS) grade and purchased from Wako Pure Chemical Industries, Ltd. (Osaka, Japan). Ultrapure water of 18.2 MΩ cm^−1^ was used throughout the experiment. For picoPPESI-MS operation, an ionic liquid, trihexyl (tetradecyl) phosphonium bis trifluoromethanesulfonyl amide (Cyphos IL109 Strem Chemicals Inc., MA, the US) was suspended in phenyl methyl silicone oil (Wacker silicone fluid AS4, Munich, Germany) at a concentration of 0.01% (v/v) to enhance electric conductivity of the silicone oil. All manipulations were conducted under a digital microscope (KH-8700, HiRox Co. Ltd., Tokyo, Japan), and kernels attached to the sample holder was humidified during all processes. In addition to the picoPPESI-MS analysis, *in situ* cell turgor in both pericarp and outer endosperm located at the same dorsal side of the kernels was independently assayed under humid conditions without removing the pericarp, as described previously (Wada et al., 2011, 2014).

### Microscopy

Kernel samples for microscopic observation were sampled in growth chambers, fixed and embedded according to the method of Saito et al. (2014) with slight modifications. Transverse segments (1–2 mm thick) from the middle of the kernel at 12, 20, and 40 DAH were fixed with 4% (w/v) paraformaldehyde in 100 mM sodium phosphate (pH 7.2) for 3 h at room temperature and then washed in 100 mM phosphate buffer (pH 7.2). Fixed tissues were dehydrated through an ethanol series, and embedded in LR White resin (London Resin, Hampshire, the UK) by two-days polymerizing at 60°C. Semi-thin sections (appropriately 900 nm) for light microscopy were stained with 0.1% (w/v) Coomassie Brilliant Blue for 1h followed by potassium iodide for 1min, and ultra-thin sections (appropriately 80–100 nm) for electron microscopy were stained with lead citrate. After the staining, ultra-thin sections were observed with a transmission electron microscope (TEM; JEM-1010, JEOL Ltd., Tokyo, Japan). For the organelle arrangement image analysis, the outline of all amyloplasts, protein bodies, and other area (referred to as ‘gap’) on the light microscopic images, and the area of both PBs on TEM images were traced by using ImageJ software (US National Institutes of Health, Bethesda, MD, the US). The outline of PSV and PBII in PSV was also traced. By assuming that they were spherical with a same *r*, the ratio of volume (*V*=4/3π*r*^3^) to the area (*A*=π*r*^2^) was 4/3*r*. This value was regarded as the representative ratio to calculate *V* from *A* with the conversion, *V*= 4/3*rA*. The volume of PSV matrix was calculated as the difference between PSV and PBII volumes.

### Protein Extraction from Rice Kernels and SDS-PAGE

In each treatment, one-third of the kernel containing the dorsal side of the matured kernels, corresponding to the chalky zone of white-back rice, was removed using a razor blade. Total proteins were extracted from the kernel section in 50 mM Tris-HCl, pH 6.8, 100 mM DTT, 8 M urea, 4% (w/v) SDS, and 10% (v/v) glycerol. Proteins were separated by SDS-PAGE with 10–20% acrylamide and stained with Coomassie Brilliant Blue.

### Kernel Quality and Weight

Spikelets in each portion of a panicle were further classified according to the method of Matsuba (1991). The appearance of dehulled grains (the numbers of white-back kernels) was visually evaluated with >1.8mm thickness of grains, according to the standard evaluation method of The Ministry of Agriculture, Forestry and Fisheries of Japan (http://www.maff.go.jp/j/seisan/syoryu/kensa/pdf/genmai_kaisetsu.pdf). The dry weight of kernel samples was determined during the treatment and at harvest, as described previously (Wada et al., 2011).

### Nitrogen and Starch Content Assay

The protein content of kernels is conventionally estimated using an N-protein conversion factor, 5.95 from the N content determined by the Kjeldahl method.

### Statistical Analysis

Statistical analysis of all other data was conducted using a Tukey’s test in a general linear model (GLM) procedure in JMP software (version 12.1.0, SAS Institute Inc., Cary, NC).

## Acknowledgements

The authors thank Ms. Fujiko Komiya, Mr. Makoto Nakajima, and Mr. Keiji Miike for their help in growing the rice plants and assistance with the experiments. This work was supported by JSPS KAKENHI Grant Number 16H02533 and 17H03759. RE-B is a research member of National Council of Scientific and Technological Research (CONICET), Argentina.

## Author contributions

Hiroshi Wada, Conceptualization, Investigation, Formal analysis, Methodology, Project administration, Supervision, Writing – original draft, Writing – review and editing; Yuto Hatakeyama, Investigation, Formal analysis, Methodology, Writing – original draft, Writing – review and editing; Yayoi Onda, Investigation, Funding acquisition; Writing – review and editing; Hiroshi Nonami, Funding acquisition; Writing – review and editing; Taiken Nakashima, Investigation, Funding acquisition, Writing – review and editing; Rosa Erra-Balsells, Methodology, Writing – review and editing; Satoshi Morita, Methodology, Validation, Writing – review and editing; Kenzo Hiraoka, Methodology, Funding acquisition, Writing – review and editing; Fukuyo Tanaka, Investigation, Writing – review and editing; Hiroshi Nakano, Validation, Writing – review and editing

## Additional files

### Supplementary files

Supplementary file 1. Diagram of the side view of the on-site cell metabolomics system placed in the laboratory (A) and expanded figure of picoPPESI-MS shown in red rectangle in A (B).

Supplementary file 2. List of amino acids and carbohydrates detected, as independent ions of clusters, in outer endosperm in each treatment using picoPPESI-MS in negative ion mode.

Supplementary file 3. Calculated area of PBI (A) and PBII (B) per outer endosperm cell at 40 DAH shown in Fig. 2G, H, I. Each parameter was calculated by multiplying the number and the averaged area of each PB.

Supplementary file 4. The transverse sections of heat-treated kernels collected at 20 DAH (A). In A, black square corresponds to the area used for light microscopic observation (C-K). Area-based percentage of each organelle per cell in a part of chalky zone (traced in red in A) at 20 DAH was shown in B.

Supplementary file 5. PicoPPESI-MS negative ion mode mass spectra obtained from the pericarp cells in 26°C (C), 34°C (D), and N+34°C (E) at the same stage. Data are representative of similar experiments with 4-5 kernels in each treatment.

